# Palmitic acid overload does not promote full-blown dysfunction in SGBS adipocytes which is reflected in their secretome

**DOI:** 10.64898/2026.11.19.767488

**Authors:** Sharon Angelini, Gabriella Stifani, Gabriele La Runa, Lucia Carmela Cosenza, Giada Lodi, Rebecca Voltan, Paola Secchiero, Fabiola Castaldo, Stefania Gessi, Stefania Merighi, Mercedes Fernandez, Anne Bigot, Martin Wabitsch, Carlo Cervellati, Juana M. Sanz, Angelina Passaro, Domenico Sergi

## Abstract

**Background:** Energy excess, and particularly long-chain saturated fatty acid (SFA) overload, are instrumental in promoting obesity and hampering metabolic health. However, SFA contribution to human adipocyte metabolic and endocrine dysfunction remains elusive. Nonetheless, we have already provided evidence that a 48-hour palmitic acid (PA) exposure is not sufficient to hamper Simpson-Golabi Behmel Syndrome (SGBS) adipocytes insulin signalling.

**Objective:** This study aims at evaluating whether a 96-hour exposure to PA is able to disrupt SGBS adipocyte metabolic health and induce the release of a secretome capable of hampering insulin signalling in skeletal muscle cells.

**Method:** SGBS adipocytes were treated with either 1000 µM PA or bovine serum albumin (BSA) for 96 hours followed by the assessment of the NFκB signalling and the expression of genes associate with inflammation and adipocyte metabolic stress. Furthermore, adipocytes were labelled with BODIPY and lipid droplet as well as cell size semi-quantified using ImageJ. Insulin-induced phosphorylation of Akt and Akt substrate of 160 kDa (AS160) was assessed by Western Blot. The effect of PA on SGBS secretome was assessed by both ELISA as well as MILLIPLEX®. Finally, human immortalised myotubes were co-cultured with BSA or PA-treated SGBS adipocytes for 24 hours followed by the assessment of insulin signalling at the Akt level.

**Results:** PA did not activate the NFκB signalling pathway, but induced the expression of *MCP-1* while tending to upregulate *IL-6* mRNA levels. Despite 96-hour PA exposure leading to an increase in lipid droplet and cell size, this was not coupled with an impairment in insulin signalling. Nevertheless, this SFA promoted an increase in IL-6 and IL-8 secretion while tending to enhance the release of adiponectin by SGBS adipocytes. Finally, the secretome from PA-treated SGBS adipocyte did not impair insulin-induced phosphorylation of Akt relative to controls in myotubes.

**Conclusions:** Thus, in the absence of the stromal vascular fraction, direct exposure of SGBS adipocytes to PA overload is not sufficient to elicit overt dysfunction. The absence of full-blown adipocyte dysfunction is reflected in the secretome of these cells which failed to disrupt skeletal muscle insulin signalling.

## 1. Introduction

The prevalence of obesity is increasing at an alarming rate worldwide ^1^, posing a significant threat to people’s health and economic burden to the healthcare systems. Indeed, obesity not only imparts a social disadvantage, hampers economic productivities ^2^ and heavily impacts on healthcare medical expenditure ^3^ but also contributes to a plethora of comorbidities, ranging from type 2 diabetes and cardiovascular disease to several types of cancer ^4–6^. In light of this, obesity must not be deemed as a mere accumulation of excess adiposity. Instead, the pathological overexpansion of the white adipose tissue, aimed at accommodating excess substrate availability, drives the dysmetabolic derangements which underpin obesity comorbidities. It translates that adipose tissue overexpansion and its consequent dysfunction are at the interface between obesity and its cardiometabolic aberrations, particularly insulin resistance ^7, 8^. Adipose tissue dysfunction is marked by insulin resistance, metabolic inflammation, characterised by the infiltration of immune cells and particularly macrophage which acquire and M1 polarisation, as well as impaired expandability ^9^. These immunometabolic defects contribute to systemic insulin resistance which fosters aberrant lipolysis. The resulting lipid spillovers promote lipotoxicity ^10^, which, in turn, is a key pathogenetic mechanism that disrupts systemic insulin sensitivity ^11^. To a similar extend adipose tissue metabolic inflammation results in the secretion of pro-inflammatory mediators, such as TNF-α and IL-6 which are able to disrupt insulin signal transduction pathway ^12–14^. However, apart from affecting the release of free-fatty acids and pro-inflammatory mediators, the dysfunction of the adipose tissue is also characterised by a disruption of its endocrine function marked by the upregulation of adipokines that foster insulin resistance, like resistin, and a concurrent downregulation of insulin sensitising hormones such as adiponectin ^15^. Additionally, it must be acknowledged that a key component of adipose tissue secretome is represented by extracellular vesicles (EVs) ^16^ able to convey proteins, lipids and nucleic acid, such as microRNAs (miRNAs) to metabolically active tissues thereby shaping their insulin sensitivity and energy metabolism ^17, 18^. In line with the disruption of the adipose tissue secretome, also the number of adipose tissue-derived EVs and their cargo is affected by obesity ^19–21^. Most importantly, the increase in adipose tissue-derived EVs and a shift in their protein, lipid and miRNA cargo are tightly linked to insulin resistance ^20, 22–25^. Thus, the expansion and metabolic dysfunction of the adipose tissue are reflected in its secretome which, in turn, contribute to systemic insulin resistance.

In this context, the skeletal muscle represents a key target of adipose tissue-derived cues in order to modulate systemic metabolic homeostasis. Indeed, the skeletal muscle is pivotal in governing glucose metabolism, as it absorbs up to 80% of the postprandial circulating levels of this metabolic fuel. Moreover, disruption of skeletal muscle insulin sensitivity represents a key defect in the pathogenesis of type 2 diabetes mellitus (T2DM) ^26, 27^. It translates that a shift towards a pro-insulin resistance adipose tissue secretome, as it occurs in the context of obesity, disrupts the adipose tissue-skeletal muscle axis paving the way for the development of insulin resistance and impairing systemic glucose homeostasis.

Along with the stromal vascular fraction, mature adipocytes also contribute to adipose tissue secretome. In particular, adipocytes have been proposed as a key contributors to adipose tissue-secreted extracellular vesicles ^28^ which, in turn, are a pivotal constituent of the secretome of fat tissue ^16^. However, whether saturated fatty acid overload directed to isolated adipocytes is sufficient to alter the metabolic function of their secretome remains to be fully elucidated. Additionally, we have previously reported that a 48-hour palmitic acid (PA) challenge was not sufficient to disrupt Simpson-Golabi Behmel Syndrome (SGBS) adipocyte metabolic health ^29^. Thus, the aim of this study is two-fold: firstly, to evaluate whether a longer PA exposure is able to disrupt SGBS adipocyte insulin signalling and promote metabolic inflammation; secondly, to assess whether the secretome released by adipocytes challenged with an energy excess is able to hamper insulin signalling in skeletal muscle cells.

## 2. Materials and methods

### 2.1. Cells culture

#### 2.1.1 SGBS human adipocytes

Human SGBS adipocytes were kindly provided by Professor Martin Wabitsch (University Hospital of Ulm, Germany) ^30^. SGBS adipocytes were cultured in Dulbecco’s modified Eagle’s medium/nutrient mixture F-12 (DMEM/F12) supplemented with 10% foetal bovine serum (FBS), pantothenic acid (17 µM), biotin (33 µM), penicillin (100 U/mL), and streptomycin (100 µg/mL), until 80%-90% confluence was reached. To induce their differentiation into mature adipocytes, a 14-day differentiation protocol involving two distinct media was applied. For the first 4 days, cells were cultured in Quick medium consisting of low glucose (5.5 mM) DMEM supplemented with pantothenic acid (17 µM), biotin (33 µM), penicillin (100 U/mL), and streptomycin (100 µg/mL), transferrin (0.01 mg/mL), insulin (20 nM), cortisol (100 nM), triiodothyronine (0.2 nM), dexamethasone (25 nM), isobutyl methylxanthine (250 µM) and rosiglitazone (2 mM). For the remaining 10 days, cells were maintained in 3FC medium, which contained the same components as the Quick medium except for the omission of dexamethasone, isobutyl methylxanthine and rosiglitazone. Cells were maintained in a humidified incubator at 37°C in a 5% CO2 atmosphere.

#### 2.1.2 AB1079 human myotubes

Immortalized human myoblast (AB1079 cell line), isolated from the quadriceps of a 38-year-old male donor, were employed as the in vitro model for skeletal muscle experiments. This cell line was kindly provided by Anne Bigot (Institute of Myology, Paris, France). Myoblasts were maintained in growth medium consisting of a 4:1 (v/v) mixture of low-glucose (5.5 mM) DMEM and M199. This formulation was supplemented with 20% FBS, penicillin (100 U/mL), streptomycin (100 µg/mL), fetuin (25 ng/mL), epidermal growth factor (5 ng/mL), dexamethasone (0.2 ng/mL) and basic fibroblast growth factor (0.5 ng/mL). Upon reaching 80% confluence, myogenic differentiation was induced. The differentiation medium consisted of low-glucose DMEM supplemented with 2% horse serum (HS), penicillin (100 U/mL) and streptomycin (100 µg/mL). Successful differentiation into mature, multinucleated myotubes was monitored and confirmed morphologically.

### 2.2 BSA-palmitic acid conjugation

Due to its lipophilic nature and poor aqueous solubility, PA was conjugated to BSA to ensure its bioavailability and mimic the physiological transport of fatty acids in the bloodstream, as previously described ^31^. Briefly, PA was dissolved in 0.1 M NaOH at 70°C using a block heater to yield a 20 mM solution. In parallel, low-endotoxin, fatty acid-free BSA was dissolved in serum and antibiotic free low-glucose DMEM to a final concentration of 0.5 mM. Once the PA was completely dissolved, it was diluted 10-fold into the BSA solution to achieve a 4:1 (PA:BSA) molar ratio (2 mM PA: 0.5 mM BSA). The mixture was then vortexed for 10 seconds and incubated at 55°C for 10 minutes to facilitate conjugation. The vehicle control (BSA) was prepared using an identical protocol, substituting the PA solution with 0.1 M NaOH. Both the PA-BSA complex and the control solution were cooled to room temperature, filter-sterilized and stored at −20°C until experimental use.

### 2.3 Adipocytes treatment

Following complete differentiation, mature adipocytes were treated for 96 h in low-glucose (5.5 mM) DMEM containing either low-endotoxin bovine serum albumin (BSA, 250 µM) as a vehicle control, or PA (1000 µM). Media containing the treatment was refreshed every 48 hours.

### 2.4 LDH cytotoxicity assay

SGBS adipocyte membrane integrity was assessed by measuring lactate dehydrogenase (LDH) release into the conditioned media using the LHD assay (Merck – Darmstadt, Germany) according to the manufacturer’s protocol. One-hundred μL of conditioned media was transferred into a 96-well plate along with 100 μL of reaction mixture. After a 30-min incubation at room temperature, the reaction was quenched by adding 50 μL of stop solution. Absorbance was subsequently measured at 500 nm using a Tecan Infinite® 200 PRO microplate reader (Tecan Trading AG, Switzerland).

### 2.5 Adipocyte and myotube co-culture

To assess adipocyte-myotube crosstalk, SGBS adipocytes were cultured and differentiated on transwell (Corning® Transwell® 24 well plates) inserts with 0.4 μm pore size membranes and exposed to PA for 96 h, as already described. After the treatment, adipocyte-containing inserts were placed over differentiated human myotubes, allowing indirect cell-to-cell communication for 24 hours.

### 2.6 Cell staining and fluorescence microscopy

To evaluate neutral lipid accumulation within lipid droplets, SGBS adipocytes were labelled using the BODIPY fluorescent probe (Thermo Fisher Scientific - Waltham, MA, USA). Following 96 h of treatment treatment with either BSA or 1000 µM PA, the culture medium was aspirated and cells were rinsed with PBS. Cells were then fixed with 4% paraformaldehyde for 15 min at room temperature. After two washes with PBS, the fixed cells were incubated with 1 µM BODIPY in PBS for 15 min in the dark. To eliminate excess fluorophore, two additional PBS washes were performed. Finally, fluorescence images were captured using an EVOS M5000 fluorescence microscope (Thermo Fisher Scientific - Waltham, MA, USA).

### 2.7 Assessment of lipid droplet number and dimensions

Following the 96 h treatment, adipocytes and lipid droplets dimensions were evaluated using ImageJ software. For each experimental group, 5 representative images were analysed; to ensure data objectivity and minimize inter-observer bias, 5 independent measurements per image were performed by 5 distinct operators. Images were scale-calibrated, converted to 8-bit grayscale, and processed to optimize contrast and reduce background noise. Overall adipocyte area was measured to quantify cell size. Intracellular lipid droplets were segmented using automated thresholding and watershed separation. Particle analysis was performed to determine droplet count and diameter, filtering out artifacts based on shape and size criteria. Finally, the total number of lipid droplets was normalized to the respective adipocyte cell area.

### 2.8 Western blot

After 96-h treatment, cells were washed with PBS and lysed with RIPA buffer supplemented with protease and phosphatase cocktail inhibitors. To assess insulin signal transduction pathway, cells were incubated with 100 nM insulin (Merck – Darmstadt, Germany) for 15 min prior lysis. Next, cell lysates were centrifuged at 14000g for 15 min to remove cells debris and proteins concentration was assessed using BCA colorimetric assay (Merck – Darmstadt, Germany). Five micrograms of protein were resolved using a 6% or 12% polyacrylamide gel and then transferred onto a polyvinylidene fluoride (PVDF) membrane using the Trans-Blot Turbo transfer system (Bio-Rad Laboratories – Hercules, CA, USA). Non-specific binding sites were blocked for 1 hour at room temperature in PBS supplemented with 0.1% Tween-20 (PBS-T) and 5% non-fat dry milk. After blocking, membranes were probed overnight at 4°C with primary antibodies: anti-pAkt/PKB (Ser473), anti-Akt/PKB, pAS160 (Thr642), anti-AS160, anti-pHSL (Ser660), anti-HSL and anti-IκBα. Consecutively, the excess of primary antibody was washed away by three 5-min washes with PBS-T and probed with HRP-conjugated secondary antibody (1:10000) for 1 h and 15 min at room temperature and then washed again with PBS-T as described above. Protein bands were revealed using Westar εC Ultra 2.0 ECL substrate (Cyanagen Srl, Bologna, Italy) and captured using Invitrogen iBright Imagers (Thermo Fisher Scientific, USA), while densitometry analysis was performed using ImageJ.

### 2.9 RNA extraction, cDNA synthesis and qPCR analysis

After the treatments, cells were lysed using the Aurum Total RNA Lysis Solution (Bio-Rad Laboratories, USA) and stored at −80°C until use. RNA was extracted using the Aurum Total RNA Mini Kit DNA-free (Bio-Rad Laboratories, USA), after which 500 ng was retro-transcribed into cDNA using 5× PrimeScript RT Master Mix (TaKaRa, Japan) according to the manufacturer’s instructions. The expression of *interleukin-6 (IL-6)*, *interleukin-1β (IL-1β)*, *fatty acid binding protein-4 (FABP-4)*, *monocyte chemoattractant protein-1 (MCP-1)*, *cell death-inducing DFFA-like effector C (CIDEC)* and *peroxisome proliferator-activated receptor gamma (PPARγ)* was assessed by the StepOne Plus Real-Time PCR system and using the TB Green Premix Ex Taq (Tli RNaseH Plus) (TaKaRa) master mix. The PCR cycles were set as follow: 1 cycle at 95 °C for 30 s, followed by 40 cycles at 95 °C for 5 s and 30 s of the annealing phase, whose temperature was primer set-specific as detailed in Table 1. Data were normalized to *18S rRNA* and analysed using the comparative ΔCt method.

### 2.10 Adiponectin quantification in conditioned media

CM were collected after 96 hours treatment, centrifuged at 1500g per 20 min to remove cell debris and stored at −20°C until use. Adiponectin secretion into the CM was assessed using ELISA kits from Abcam (Cambridge, UK) following the manufacturer’s instructions.

### 2.11 Cytokine multiplex assay

The quantitative measurement of proinflammatory mediators released in adipocyte supernatant was conducted by utilizing the MILLIPLEX® Human Cytokine/Chemokine/Growth Factor Panel A (catalogue number HCYTA-60k, Merck Millipore, Darmstadt, Germany). The panel can perform simultaneous quantification of the following 38 human proteins. Frozen samples were thawed on ice and analysed in accordance with the manufacturer’s instructions. In summary, undiluted samples and kit standards were subjected to an overnight incubation at 4°C with antibody-conjugated beads within the provided 96-well plate. Subsequently, detection antibodies were added and the mixture was left to incubate for 1 hour at room temperature. This was followed by the addition of streptavidin-phycoerythrin and a further 30 minutes of incubation. Following three additional washes, the beads were resuspended in Drive Fluid and analysed using a Luminex 200 system (Luminex, Austin, TX, USA) with xPONENT Acquisition and MILLIPLEX Analyst 5.1 software (Merck, Darmstadt, Germany). Protein concentrations were calculated by interpolation from standard curves and expressed as log transformation of pg/mL.

### 2. 12 Data analysis

Statistical analyses were performed using GraphPad Prism version 8.0.1 for Windows (GraphPad Software Boston, Massachusetts USA, www.graphpad.com). Data are expressed as mean of at least 3 biological replicates ± standard error mean (SEM) and differences between treatments were performed using Student’s T-test, one-way or two-way ANOVA procedure followed by Tukey’s multiple comparison. A *p – value* ≤ 0.05 was considered significant.

## 3. Results

### 3.1 Palmitic acid induces SGBS adipocyte hypertrophy in the absence of cytotoxicity

PA has previously been shown to promote adipocyte hypertrophy, albeit in murine adipocyte models ^32–34^. This was confirmed in human SGBS adipocytes, with this long-chain saturated fatty acid inducing an increase in lipid droplet area (p< 0.01) **(Figure 1A & B)** which was paralleled by a concomitant increase in cell area (p< 0.01) **(Figure 1A & C)** and a trend towards an upregulation of PLIN1 (p= 0.056) **(Figure 1D & E)**. However, these effects occurred in the absence of an increase in the number of lipid droplets per cell (p> 0.05) **(Figure 1A & F)** and cytotoxicity as indicated by the fact that PA did not increase lactate dehydrogenase activity in the cell culture media (p> 0.05) **(Supplementary figure 1)**. Additionally, PA was readily taken up by the adipocytes and stored as indicated by the increase in the fluorescent signal within lipid droplets upon exposure to a fluorescently-labelled PA analogue **(Figure 1G).**

**Figure 1.**
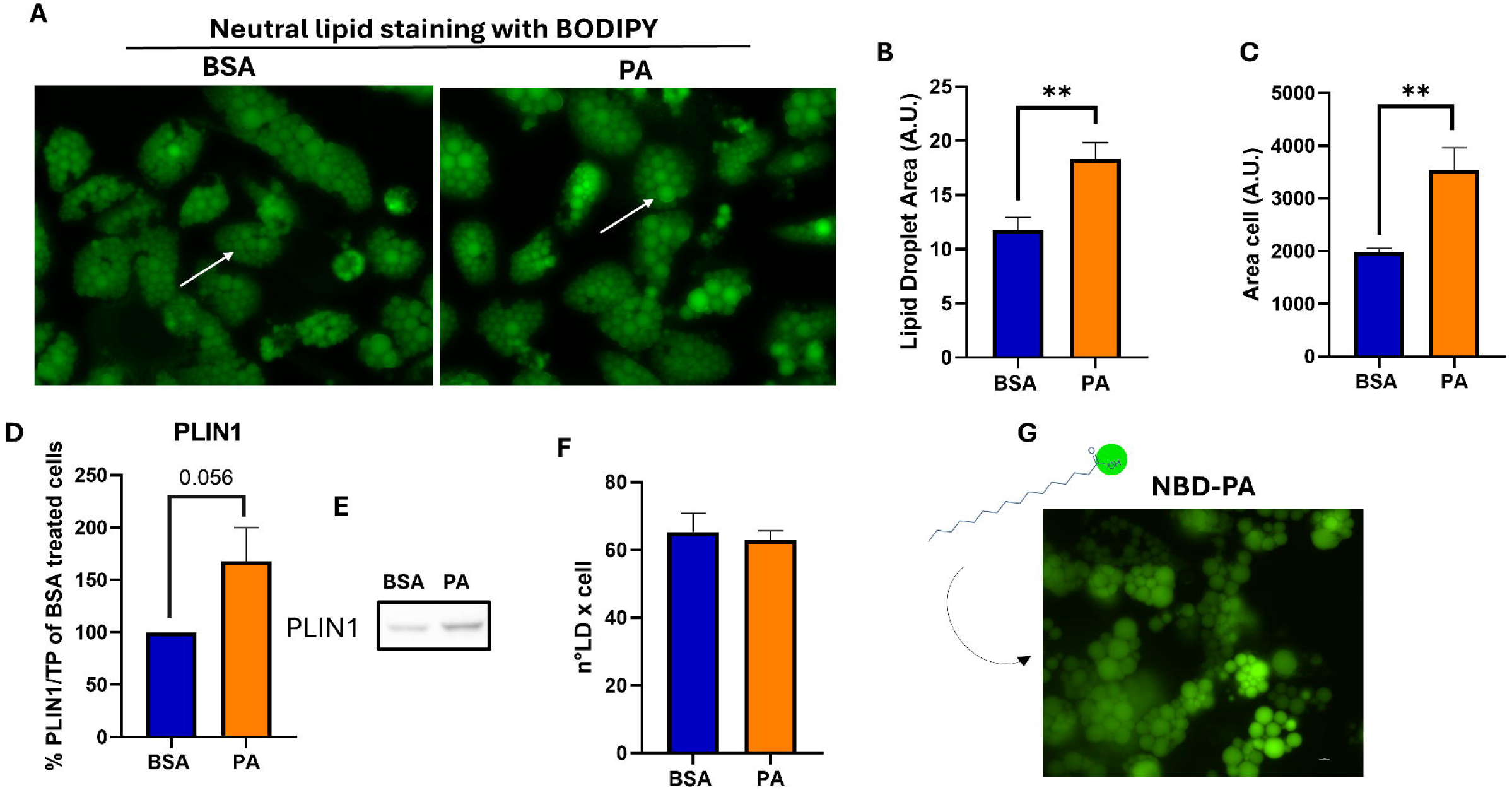
Effects of PA on lipid droplet size. (A) Fluorescence microscopy of BSA and PA-treated adipocytes stained with Bodipy neutral lipid dye. Measures of (B) lipid droplet area, (C) adipocyte area and (C) lipid droplet count obtained using ImageJ software. (E) Densitometric analysis of PLIN1 expressed as percentage of BSA-treated cell and (F) its representative western blot. BSA, Bovine albumin serum; PA, Palmitic acid; NBD-PA, [16-[(7-nitro-2-1,3-benzoxadiazol-4-yl)amino] Palmitic acid; LD, Lipid droplets. ** p – value < 0.01.

### 3.2 SGBS adipocytes displayed early signs of metabolic stress in the absence of overt metabolic dysfunction in response to PA

Considering an increase in adipocyte size following energy overload representing a pivotal trigger for the onset of adipose tissue dysfunction ^8^, SGBS adipocytes were profiled further to evaluate if this long-chain saturated fatty acid induced the expression of surrogates of adipocyte metabolic distress. FABP4 is pivotal in regulating adipocyte size and links adipocyte dysfunction with systemic metabolic health ^35^. As such, its expression was assessed in response to PA challenge to evaluate whether this saturated fatty acid induced early signs of metabolic deterioration which, in turn, may rebound upon systemic metabolic health. PA significantly upregulated FABP4 (p <0.001) **(Figure 2A)**. To further evaluate whether this long-chain saturated fatty acid hampered adipocyte function, and particularly storage capacity, it was assessed the expression of *CIDEC* given its role in regulating adipocyte lipid handling and incorporation into lipid droplets ^36, 37^. Contrarily to its effect of *FABP4*, PA did not affect the expression of *CIDEC* (p> 0.05) **(Figure 2B)**. Furthermore, *peroxisome proliferator-activated receptor γ* (*PPARγ*) is a pivotal regulator of adipocyte and systemic metabolic health ^38^, therefore its expression was assessed to gain further insights on the metabolic effects of PA overload. In line with the effect on *CIDEC*, PA did not modulate the expression of *PPARγ* (p> 0.05) **(Figure 2C)**.

**Figure 2.**
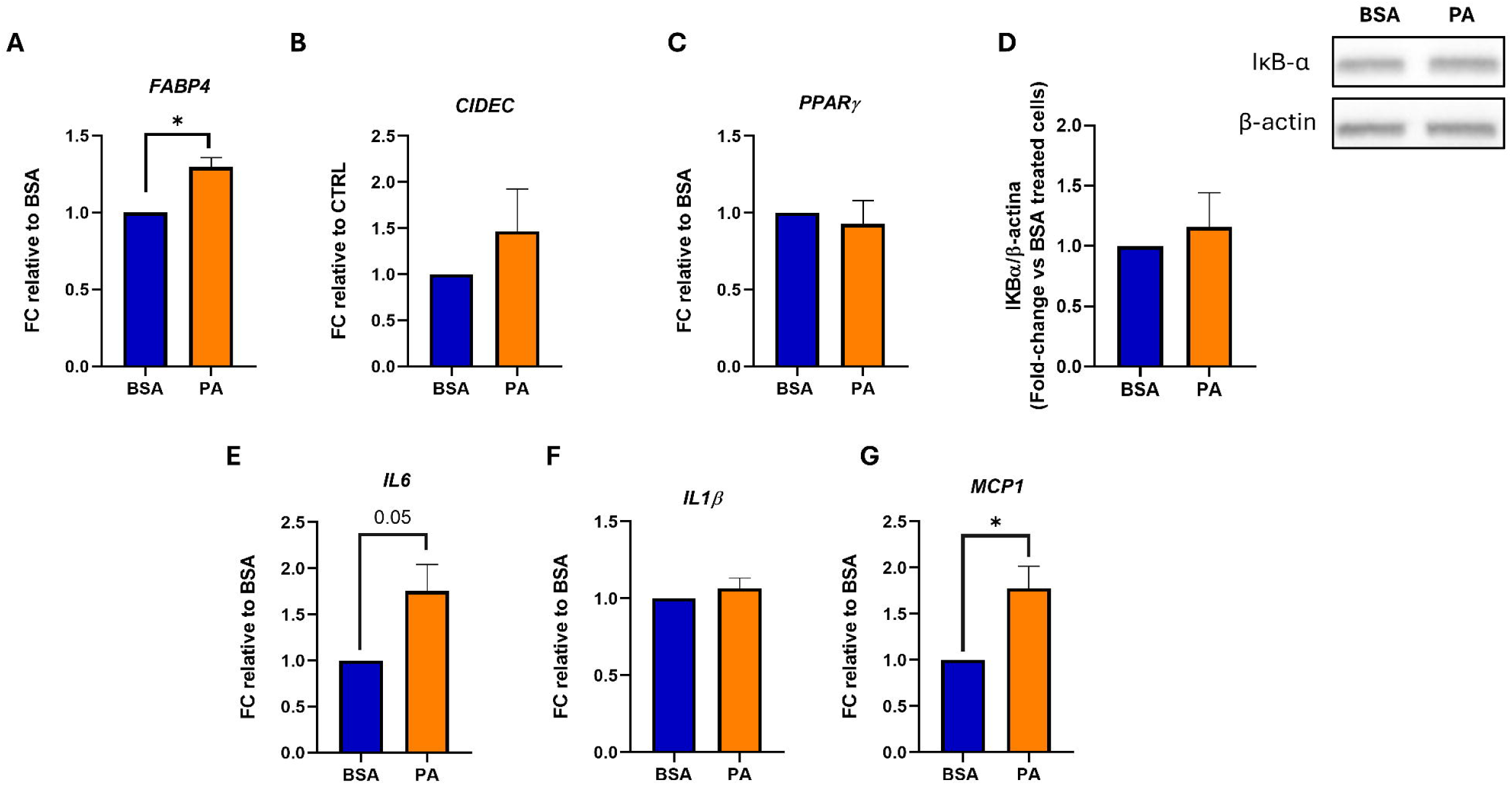
Effect of PA on markers of adipocytes dysfunction. Gene expression analysis of (A) *FABP4*, (B) *CIDEC*, (C) *PPARγ*, (D) *IL6*, (E) *IL1β* and (G) *MCP1*. (D) Densitometric analysis and representative blot of IκBα normalised over β-actin and expressed as fold-change versus BSA-treated cells. BSA, Bovine albumin serum; PA, Palmitic acid; FABP, Fatty Acid-Binding Protein 4; CIDEC, Cell Death Inducing DFFA Like Effector C; PPARγ, Peroxisome Proliferator-Activated Receptor Gamma; IL6, Interleukin 6; IL1β, Interleukin 1 beta; MCP1, Monocyte chemoattractant protein 1; IκBα, Inhibitor of nuclear factor kappa B alpha. * p – value ≤ 0.05

The activation of metabolic inflammation is a hallmark of lipotoxicity ^39, 40^ as well as adipose tissue dysfunction ^41^, therefore it was assessed if PA overload triggered the activation of inflammatory pathways. The 96-hour exposure to PA did not activate the NFκB signalling pathway as indicated by the inability of this fatty acid to decrease the abundance of inhibitor of NF-κB (IKBA) (p> 0.05) **(Figure 2D)**. However, PA tended to upregulate the mRNA levels of *IL-6* (p= 0.05) **(Figure 2E)** without affecting the expression of *IL-1β* **(Figure 2F)**. Finally, PA induced *MCP-1* expression (p< 0.05) **(Figure 2G)** which further supports the early stress response mounted by adipocyte in response to energy overload.

### 3.3 Palmitic acid did not cause insulin resistance in SGBS adipocytes

To evaluate whether adipocyte hypertrophy and upregulation of specific inflammatory mediators triggered by PA was coupled with the onset of insulin resistance, insulin signalling and its ability to inhibit HSL were evaluated. PA did not affect insulin-induced phosphorylation of Akt **(Figure 3A & C)** and its substrate AS160 **(Figure 3B &C)** (all p> 0.05). In further support of this, PA did not impair the ability of insulin to inhibit the phosphorylation of HSL (p> 0.05) **(Figure 3D & E).** However, this long-chain saturated fatty acid increased the basal phosphorylation state of HSL (p< 0.05) **(Figure 3D & E)**.

**Figure 3.**
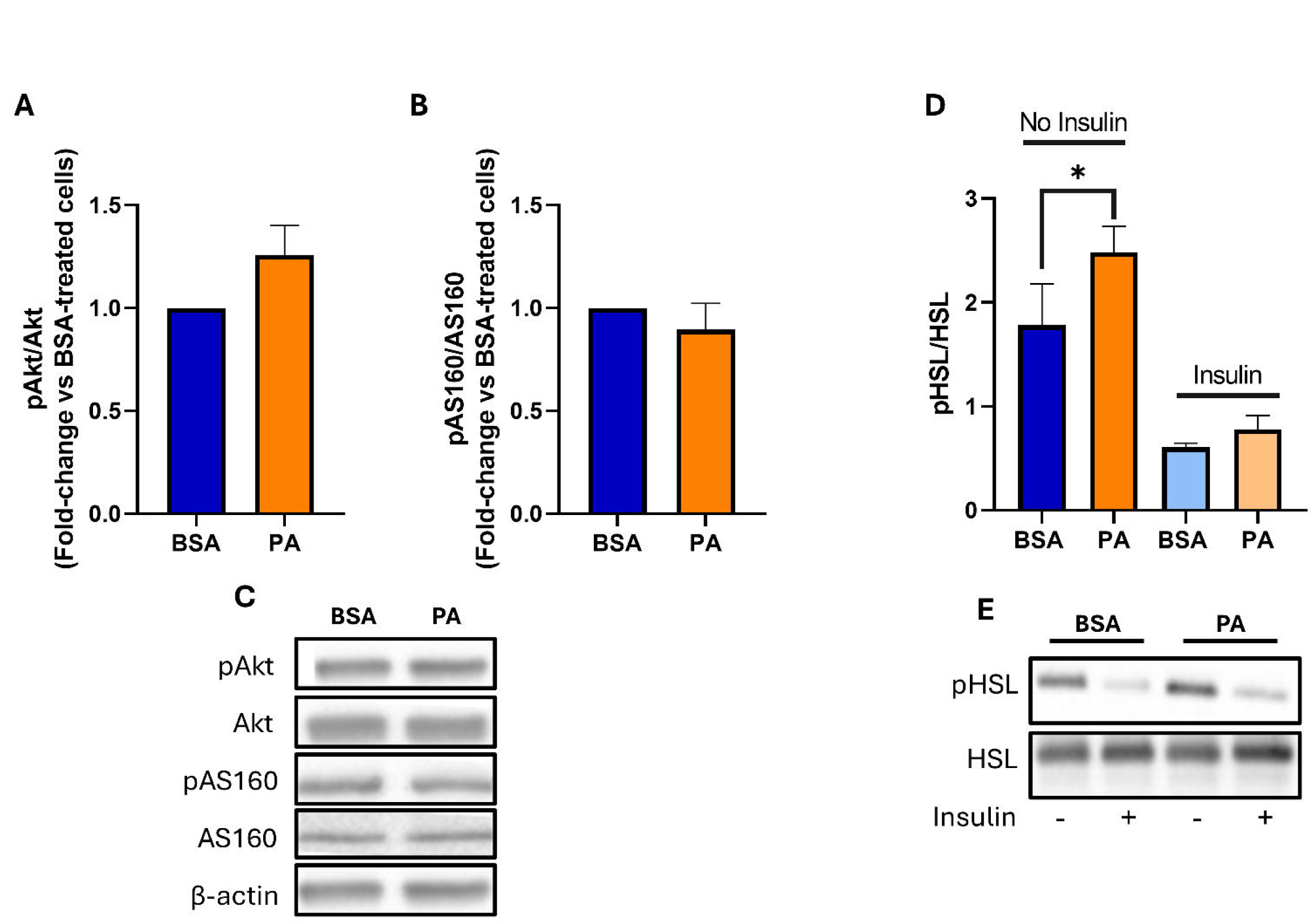
The impact of PA on insulin-mediated pathways. Densitometric analysis of insulin-mediated phosphorylation and activation of (A) Akt (Ser473), (B) AS160 and its representative Western Blots. Data were normalised over β-actin and expressed as fold-change versus BSA-treated cells. (C) Densitometric analysis of basal and insulin-mediated phosphorylation and inhibition of HSL normalised over total protein. BSA, Bovine albumin serum; PA, Palmitic acid; Akt, Protein kinase B; AS160, Protein kinase B substrate of 160 kDa; HSL, Hormone sensitive lipase. * p – value ≤ 0.05

### 3.4 Palmitic acid exposure partially altered the secretome of SGBS adipocytes which did not hamper insulin signalling recipient myotubes

The adipocyte secretome was characterised using a multiplex assay with high sensitivity, enabling the quantitative assessment of multiple soluble mediators from minimal sample volumes simultaneously. Given that energy excess and the subsequent increase in adipocyte energy stores is able to modulate the secretome of fat cells, it was assessed whether PA overload affected the secretome of SGBS adipocytes. The exposure of adipocytes to PA for 96 hours significantly increased the secretion of IL-6 and IL-8 (p< 0.05) and tended to increase the release of eotaxin (p= 0.074) **(Figure 4A).** However, contrarily to this mild increase in inflammatory mediators, PA tended to upregulate the secretion of the insulin sensitising hormone adiponectin (p= 0.067) **(Figure 4B)**.

**Figure 4.**
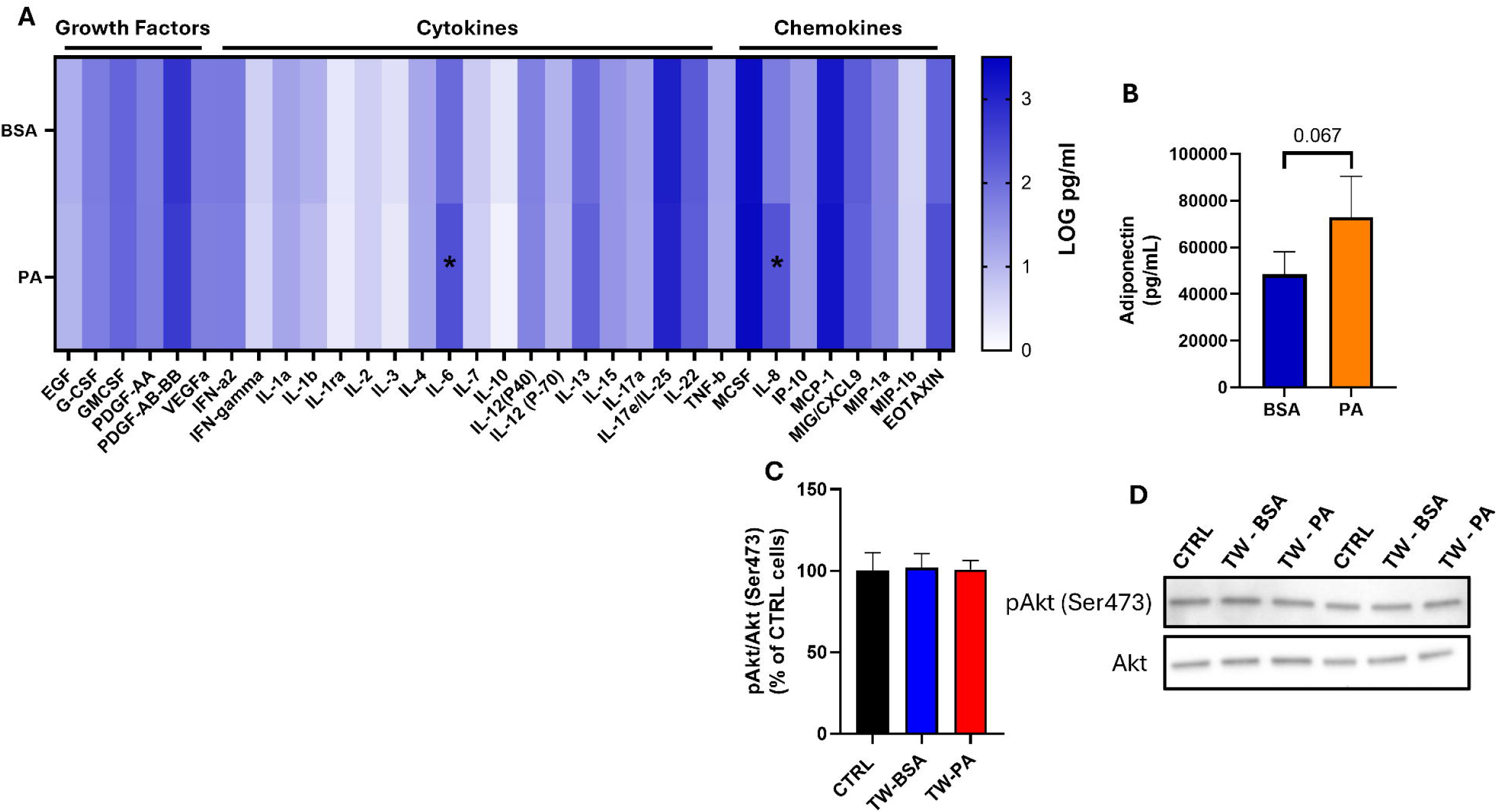
PA affects adipocytes secretome but not its effect on skeletal muscle insulin signalling. (A) Heat map representing 38 human proteins quantified via MILLIPLEX® analysis and expressed as log-transformed of pg/mL. (B) Adiponectin levels (pg/mL) quantified via ELISA assay. (C) Representative western blots and (D) its densitometric analysis of of insulin-mediated phosphorylation and activation of Akt (Ser473) of myotubes co-cultured with BSA- and PA-treated adipocytes and its relative control (CTRL) without adipocytes. * p – value ≤ 0.05; TW, transwell co-culture.

The fact that adipocytes maintain an intact insulin signalling despite being exposed to saturated fatty acid overload does not necessarily mean their secretome does not impact other metabolically active tissues. Thus, to functionally characterise the SGBS adipocyte secretome, these cells were co-cultured with immortalised human myotubes. In line with the mild effect elicited by saturated fatty acid exposure, the secretome of SGBS adipocytes treated with PA, compared to BSA, did not impair insulin-induced phosphorylation of AKT (Ser473) (p> 0.05) **(Figure 4C & D)**.

## Discussion

The results of the present study provide support to our previous findings ^29^, in that SGBS adipocytes have a remarkable capacity to store excess long-chain saturated fatty acids within lipid droplets. However, after a 96-hour PA challenge, neutral lipid accumulation within lipid droplets led to adipocyte hypertrophy coupled with the onset of early sign of metabolic stress, albeit in the absence of overt adipocyte dysfunction and insulin resistance. Finally, despite PA increased the secretion of some pro-inflammatory mediators by SGBS adipocytes, the secretome of these cells did not hamper insulin signalling in recipient myotubes.

The increase in lipid droplet size observed following 96-hour PA exposure suggests that the excess of this long-chain saturated fatty acid is effectively taken-up by SGBS adipocytes and stored in the form of neutral lipids as indicated by BODIPY staining. Even though, triglycerides were not directly measured, it is likely that they represent the main form under which PA overload is stored. Moreover, the induction of PLIN-1 abundance confirms that SGBS adipocytes orchestrate a metabolic response to accommodate energy excess in lipid droplets and protect their triglyceride content ^42, 43^. The preserved adipocyte storage capacity was further corroborated by the inability of PA to downregulated the expression of CIDEC, considering its role in lipid handling and accumulation into lipid droplets ^36, 37^. While storing excess energy within lipid droplets is pivotal in preventing the metabolically detrimental effects of lipotoxicity ^9, 44^, lipid droplet and adipocyte storage capacity can be overwhelmed by excess nutrient supply ^45^. Despite storing excess energy within lipid droplets being an adaptive mechanism to prevent lipotoxicity, chronic fuel overload is instrumental to promote adipocyte hypertrophy causing these cells to reach their critical storage threshold capacity. In this regard, although in the SGBS adipocytes did not display overt dysfunction, the signs of immunometabolic deregulation elicited by 96-hour PA exposure, albeit mild, may mark the initial stages of metabolic deterioration and indicate that adipocytes are approaching their critical storage threshold. Adipocyte metabolic stress elicited by PA was marked by the upregulation of *FABP4* and *MCP*-*1*, with *IL-6* also tending to be upregulated in response to this long-chain saturated fatty acid. FABP4, a protein highly expressed in adipocytes ^46^, has been proposed to bind free fatty acids within the cytoplasm during lipolysis to prevent their inhibitory effects on hormone sensitive lipase ^47^. The upregulation of FABP4 may be a direct consequence of increased fatty acid bioavailability in the form of exogenous PA and possibly increased basal lipolysis, given that this saturated fatty acid also increase HSL phosphorylation. The fact that the upregulation of FABP4 may be synonymous with adipocyte early metabolic deterioration is in agreement with its dysmetabolic effects ^35^. In line with this, mice knock-out for FABP4 did not develop insulin resistance despite the onset of diet-induced obesity ^48^. Additionally, an increase in FABP4 has been associated with insulin resistance as well as the development of the metabolic syndrome and T2DM ^49–51^. However, it remains to be confirmed whether the upregulation of *FABP4* in response to PA also results in an increase in its secretion.

The fact that 96 hour PA triggers a stress response is also corroborated by the upregulation of *MCP-1 and* the increase in *IL-6* mRNA levels which, however, did not reach statistical significance. The upregulation of these inflammatory mediators is in line with the pro-inflammatory effects of PA on murine adipocytes ^52, 53^. However, in SGBS adipocytes a 48 hour exposure to PA was not sufficient to modulate either *MCP-1* or *IL-6*. This suggests a longer exposure to this long-chain saturated fatty acid having caused adipocyte to approach a critical storage threshold thereby initiating a metabolic stress response. In particular, the upregulation of *MCP-1* may represent a stress signal adipocytes call upon in order to recruit macrophages ^54, 55^ which, *in vivo*, infiltrate the adipose tissue to fuel metabolic inflammation and insulin resistance ^55, 56^. However, while the induction of *MCP-1* expression may indicate the onset of a metabolic stress response, this upregulation was not corroborated by an increase in the secretion of its protein following the PA challenge. On the contrary, the trend towards an upregulation in *IL-6* expression was consistent with an increase of the secretion of this cytokine further supporting the inflammatory response elicited by PA. Importantly, the increase in *MCP-1* expression may not be directly driven by PA overload but may be regulated by IL-6. Indeed, IL-6 has been proposed to induce *MCP-1* expression ^57^ suggesting the upregulation of this pro-inflammatory cytokine representing an early signal of metabolic distress secreted by hypertrophic adipocytes to promote macrophage infiltration of adipose tissue ^58^. Nevertheless, the upregulation of *MCP-1* and the increase in IL-6 secretion occurred in the absence of NFκB activation despite both mediators being under the control of this transcription factor ^59^. The NFκB also controls the expression of *IL-1β* ^59, 60^ whose expression was not affected by the 96-hour exposure to PA thereby corroborating with the inability of the fatty acid overload to activate the NFκB signalling pathway. Thus, the effects of PA on *MCP-1* expression and IL-6 mRNA and protein secretion may occur via the activation of transcription factors other than the NFκB which, however, was not assessed as part of this study. The inflammatory response elicited by PA is further supported by an increase in IL-8 secretion by SGBS adipocytes. However, while this pro-inflammatory chemokine has been reported to be induced in the visceral adipose tissue secretome of obese individuals, compared to their lean counterparts, its increase appear to be driven by non-fat cells which have been proposed as the primary driver of the rise in IL-8 circulating levels in obesity ^61, 62^. Nonetheless, isolated adipocytes are also able to secrete this pro-inflammatory mediator ^61–64^ with its secretion being a direct function of adipocyte size ^65^. This paradigm is in agreement with the present findings reporting an increase in both cell size and in IL-8 secretion by SGBS adipocytes which underwent PA treatment.

Despite PA triggering an inflammatory response, albeit mild, it failed to hamper insulin-induced phosphorylation of Akt and AS160. This notion was also confirmed by the ability of insulin to inhibit HSL phosphorylation to the same extent in PA and BSA-treated cells. The capacity of SGBS adipocytes to retain their insulin signalling in spite of long-chain saturated fatty acid overload may be dependent on the magnitude of the inflammatory response, which may not be sufficient to hamper insulin sensitivity. Thus, despite both IL-8 and IL-6 having been reported to impair insulin sensitivity in adipocytes ^63, 66^ and to be upregulated in the fat cells of insulin resistant individuals ^66^ their increase, albeit statistically significant, may have not been sufficient to elicit a biologically meaningful effect on insulin sensitivity. Furthermore, despite PA leading to an increase in adipocyte energy storage, this long-chain saturated fatty acid tended to increase adiponectin secretion. Adiponectin, in turn, by acting in a paracrine and/or autocrine fashion may have prevented the deterioration of insulin signalling induced by pro-inflammatory cytokines in adipocytes ^67, 68^. Furthermore, the preserved insulin signal transduction in SGBS adipocytes may be a direct function of the capacity of these cells to effectively store excess energy in the form of triglycerides within lipid droplets which is deemed as metabolically protective ^9, 44^. This, in turn, may have prevented the buildup of lipotoxic lipid species such as ceramides which, instead, are pivotal drivers of insulin resistance ^69^. However, these findings may not necessarily reflect the impact of saturated fatty acid overload on adipose tissue insulin sensitivity *in vivo*. Indeed, *in vivo,* an increase in IL-6 secretion by adipocytes, as shown herein following PA exposure, represents a pivotal trigger to recruit and promote macrophage accumulation within the adipose tissue by a noncanonical mechanism ^58^, with these immune cells being instrumental in fostering adipocyte insulin resistance ^70^. Thus, in an isolated adipocyte system, in the absence of M1-polarised macrophages, long-chain saturated fatty acid excess *per se*, may not be sufficient to promote insulin resistance.

Even though PA exposure promoted the secretion of pro-inflammatory mediators, namely IL-6 and IL-8 and potentially increased lipid spillover due to HSL basal activation, myotubes co-cultured with SGBS adipocyte-preserved their insulin signalling. In spite of the lack of direct evidence of the effect of IL-8 on skeletal muscle are lacking, this chemokine may potentially hamper insulin signalling ^63^. However, when considered in the context of the whole adipocyte secretome, it failed to hamper myotube insulin signalling. To a similar extent, the increase in IL-6 secretion by SGBS adipocytes was not paralleled by a disruption of insulin signalling in myotubes. This, in turn, may be explained by the controversial role of this cytokine when considering its effects on insulin sensitivity. In line with this, while same evidence described it as being able to cause insulin resistance, albeit its effects being modest ^58^, other reports suggest IL-6 eliciting acute insulin sensitising or no effects in healthy humans ^71, 72^. Similarly, the effects of IL-6 on insulin sensitivity are context dependent with acute exposure exerting insulin sensitising effects whereas a chronic exposure to this cytokine promoting insulin resistance ^73^. Although the increase in IL-6 and IL-8 within SGBS adipocyte secretome did not led to an impairment in myotube insulin signalling, it remains to elucidate if this effect is driven by an intrinsic inability of these cytokines to directly disrupt insulin signal transduction in myotubes or whether their metabolically detrimental role is countered by other components of the SGBS adipocyte secretome. In this regard, the induction of adiponectin secretion in response to PA, albeit not statistically significant, may putatively explain this conundrum. Indeed, adiponectin is known to exert insulin sensitising properties ^74, 75^ which may explain the preserved insulin signalling in myotubes upon exposure to adipocyte-secretome. However, the increase in adiponectin secretion following PA treatment is in disagreement with other studies which, instead, reported a downregulation of this adipokine in response to saturated fatty acid overload ^76, 77^. These discrepancies may be related to the model used to assess the effects of PA on adiponectin secretion, with previous studies describing the effects of this saturated fatty acid on murine 3T3-L1 cells ^76, 77^ as opposed to SGBS adipocytes. Despite the effects of PA on the release of adiponectin by SGBS adipocytes being counterintuitive, this may be related to the fact that at least after 96 hour treatment these cells do not display signs of overt dysfunction. Thus, adiponectin increase in the conditioned media may represent a compensatory mechanism that adipocytes adopt prior to the onset of full blown lipotoxic stress.

To conclude, PA induced a moderate metabolic distress in SGBS adipocytes, which, despite being coupled with an increase in adipocyte volume, did not suffice to hamper insulin signalling. Most importantly, this metabolic insult did not result in the release of a pro-insulin resistance secretome, with myotubes preserving their insulin signalling despite being exposed to adipocyte-derived mediators. This suggests energy surplus *per se*, when directed exclusively to adipocytes and in the absence of adipose tissue stromal vascular fraction, being insufficient to disrupt adipocyte endocrine function. Thus, the ability of SGBS adipocytes to metabolically adapt to a PA overload and the consequent absence of overt adipocyte dysfunction is reflected in the secretome of these cells which failed to disrupt skeletal muscle insulin signalling.

## Acknowledgement

This work was supported by the University of Ferrara 2024 and 2025 fund for Scientific Research (FAR) and Department of Translational Medicine, University of Ferrara, 2024 and 2025 fund to incentivize departmental research (FIRD).

## Author contributions

SA: Methodology, Formal analysis, Writing – original draft, Writing – review & editing. GS: Methodology, Formal analysis, Writing – original draft, Writing – review & editing. GLR: Methodology, Formal analysis, Writing – review & editing. LCC: Methodology, Formal analysis, Writing – review & editing. GL: Methodology, Formal analysis, Writing – review & editing. RV: Methodology, Writing – review & editing. PS: Methodology, Writing – review & editing. FC: Writing – review & editing. SG: Writing – review & editing. SM: Writing – review & editing. MF: Methodology, Writing – review & editing. AB: Methodology, Writing – review & editing. MW: Writing – review & editing. CC: Writing – review & editing. JMS: Writing – review & editing. AP: Writing – review & editing, Supervision, Writing – original draft. DS: Conceptualization, Writing – review & editing, Supervision, Writing – original draft, Funding acquisition.

## Data Availability Statement

Data are available upon request to the corresponding authors

---

**Figure S1. PA impact on cell viability.** (A) Brightfield images of (A) BSA and (B) PA-treated cells. Scale bar = 100 μm. (C) Comparison of LDH activity between BSA- and PA-treated cells expressed as Fold-Change relative to BSA-treated cells. BSA-treated cells. BSA, Bovine albumin serum; PA, Palmitic acid; LDH, Lactate dehydrogenase.

