## Supplementary figures and images for "Palmitic acid overload does not promote full-blown dysfunction in SGBS adipocytes which is reflected in their secretome"

### Supplementary Figure 1

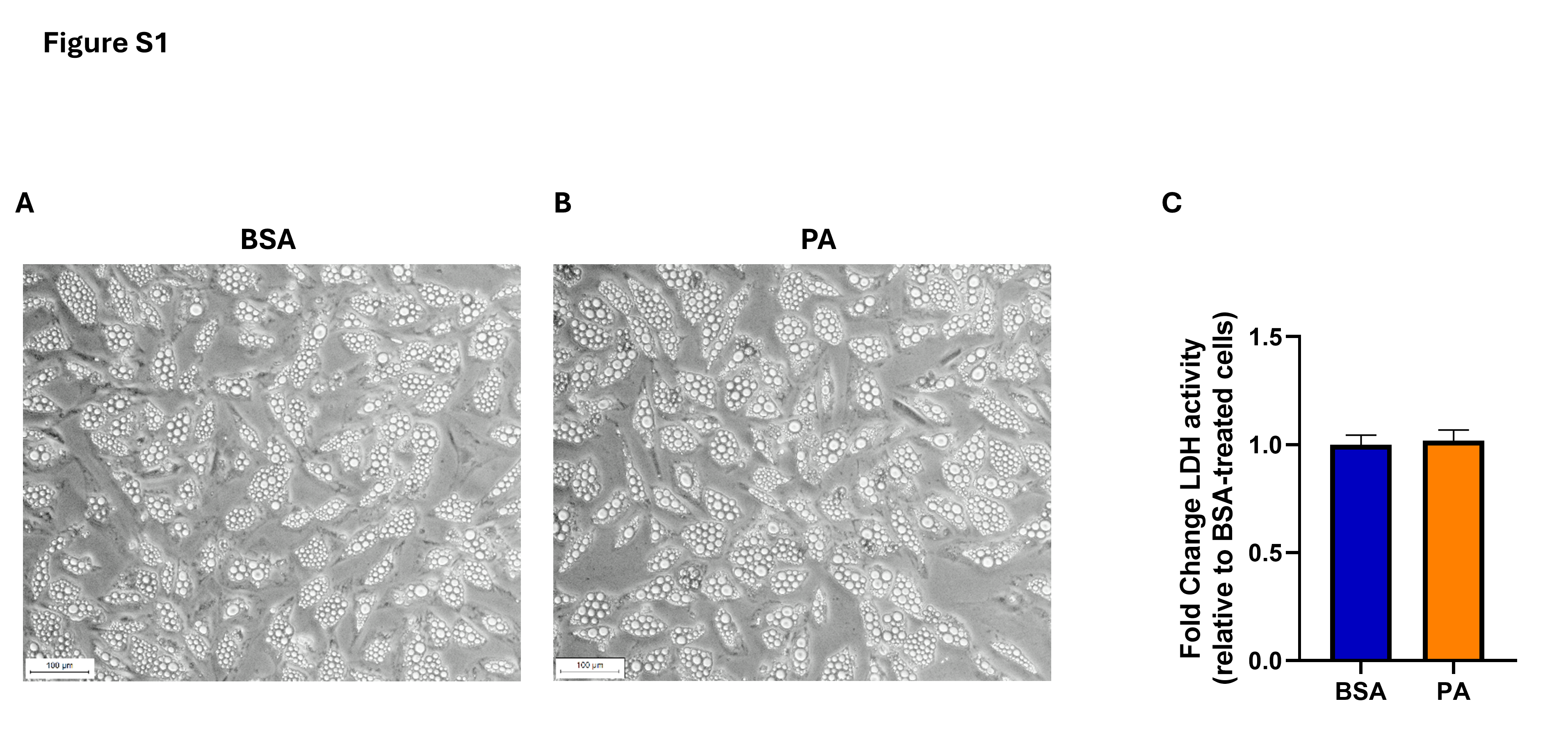
